# Visualizing endogenous RhoA activity with an improved localization-based, genetically encoded biosensor

**DOI:** 10.1101/2021.02.08.430250

**Authors:** Eike K. Mahlandt, Janine J. G. Arts, Werner J. van der Meer, Franka H. van der Linden, Simon Tol, Jaap D. van Buul, Theodorus W. J. Gadella, Joachim Goedhart

## Abstract

Rho GTPases are regulatory proteins, which orchestrate cell features such as morphology, polarity and movement. Therefore, probing Rho GTPase activity is key to understanding processes such as development, cell migration and wound healing. Localization-based reporters for active Rho GTPases are attractive probes to study Rho GTPase-mediated processes, in real time with subcellular resolution in living cells and tissue. Until now, relocation RhoA biosensors seem to only be useful in certain organisms and have not been characterized well. In this paper, we systematically examined the contribution of the fluorescent protein and RhoA binding peptides, on the performance of localization-based sensors. To test the performance, we compared relocation efficiency and specificity in cell-based assays. We identified several improved localization-based, genetically encoded, fluorescent biosensors for detecting endogenous RhoA activity. This enables a broader application of RhoA relocation biosensors, which was demonstrated by using the improved biosensor to visualize RhoA activity, during cell division, during random migration, at the Golgi membrane and induced by G protein-coupled receptor signaling. Due to the improved avidity of the new biosensors for RhoA activity, cellular processes regulated by RhoA can be better understood.

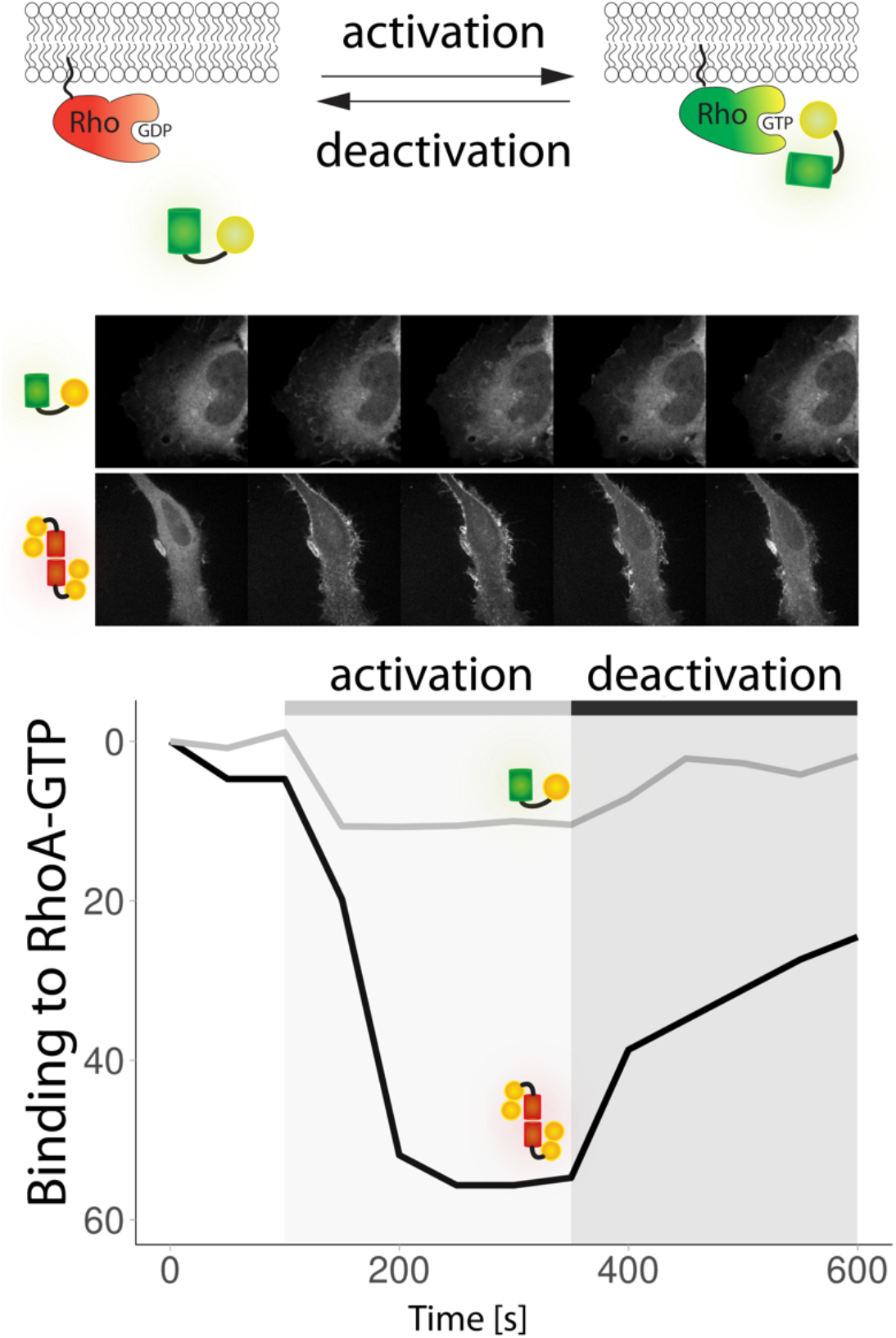

## Introduction

Rho GTPases function as a molecular switch: turned on when guanosine triphosphate (GTP) is bound and turned off when GTP is hydrolyzed to guanosine diphosphate (GDP) (Bos et al., 2009; Ridley, 2015). The conversion from inactive, GDP-bound Rho GTPase, to active, GTP-bound Rho GTPase, requires Rho guanine exchange factors (Rho GEFs) (Rossman et al., 2005). The bound GTP is hydrolyzed to GDP by the intrinsic but slow GTPase activity, thereby inactivating the Rho GTPase. This process is enhanced by GTPase-activating proteins (GAPs) (Bos et al., 2009). RhoA is one of 20 Rho GTPases and has two closely related homologues, RhoB and RhoC (Burridge & Wennerberg, 2004; Wheeler & Ridley, 2004). Although these three Rho GTPases may have different functions, we generally refer to RhoA in this manuscript. Active RhoA mainly localizes at the plasma membrane, due to its prenylated C-terminus (Garcia-Mata et al., 2011). RhoA and other Rho GTPases orchestrate the cytoskeleton dynamics and thereby cell features such as adhesion, cell migration, cell division, cell morphology and polarity (Lawson & Ridley, 2018). Therefore, they are involved in complex processes like transendothelial migration and wound healing (Heemskerk et al., 2014). Rho GTPase signaling occurs in a spatial and temporal defined manner. While biochemical assays are well established and sensitive, they only show the average of a population with no spatial resolution and a poor time resolution, in the order of minutes (Pertz, 2010). To address this issue, genetically encoded fluorescent biosensors have been engineered. These tools enable the visualization of protein activities in single, living cells with micrometer spatial and sub-second temporal resolution (Greenwald et al., 2018; Mehta & Zhang, 2011; Miyawaki & Niino, 2015).

Several genetically encoded biosensors are available to visualize active (GTP-bound) Rho GTPase. These sensors can broadly be divided in two classes, i.e. Förster resonance energy transfer (FRET)-based biosensors and localization-based biosensors (Pertz, 2004). Each class has its own advantages and disadvantages (Pertz, 2010). Regardless of the type, the sensors use a G protein binding domain (GBD), which has a higher affinity for the active GTP-bound state of the Rho GTPase relative to the inactive, GDP-bound Rho GTPase. Unimolecular Rho GTPase FRET-based biosensors consist of the Rho GTPase itself, a GBD and a FRET pair. The sequence of the domains is an important aspect of the optimization of these sensors (Fritz et al., 2013). The Dimerization Optimized Reporter for Activation (DORA) RhoA-based FRET sensor consists of RhoA and the GBD of protein kinase N1 (PKN1) (Van Unen et al., 2015). Upon Rho GTPase activation the binding domain binds the GTP-bound Rho GTPase. This conformation change leads to a FRET ratio change with a relatively small dynamic range. By design, these FRET sensors report on guanine exchange factor (GEF) activity, instead of visualizing endogenous RhoA-GTP. In contrast, localization-based sensors solely consist of a fluorescent protein fused to a GBD, which has a high affinity for the active GTP bound state. These sensors visualize endogenous RhoA-GTP. For instance, when Rho GTPase activation occurs locally at the plasma membrane, the sensor will accumulate at that location. A potential drawback is that background signal of the unbound biosensor in the cytosol, which may occlude the bound pool and reduce the dynamic range. Plus, most of the Rho GTPase binding domains are able to bind different Rho GTPases, so relocation sensors tend to be less specific than FRET sensors (Pertz, 2004). Unlike the relocation sensor, the FRET-based biosensor also contains the Rho GTPase and thereby its specificity is not only determined by the binding specificity of the GBD. For example, by changing the Rho GTPase homologue, a certain level of specificity can be achieved for RhoA, RhoB and RhoC (Reinhard et al., 2016). A key advantage of relocation probes is their simple design, utilizing only a GBD and a single fluorescent protein. Usage of a single fluorescent protein simplifies multiplexing of biosensors or the combination with optogenetics. However, the greatest advantage of the localization-based biosensor is the visualization of endogenous RhoA activity at it is unaltered location in the cell. The RhoA FRET sensors achieve subcellular resolution to a certain extent, but due to their design they do not localize as endogenous RhoA. Since temporal and spatially tightly regulated Rho GTPase activity is important for their functionality, we set out to test the RhoA relocation biosensor for its ability to visualize endogenous RhoA activity with a high spatial resolution.

Thus far, two RhoA relocation biosensors have been published. Firstly, the relocation RhoA sensor based on anillin, consisting of a GBD, C2 and pleckstrin homology (PH) domain, called anillin-homology domain (AHD)+PH-GFP or Anillin Rho binding domain (AniRBD), which was first described in 2000 (Munjal et al., 2015; Piekny & Glotzer, 2000). Secondly, the rhotekin G protein binding domain (rGBD)-based eGFP-rGBD RhoA sensor, that was reported in 2005 (Benink & Bement, 2005). Different versions appeared over the years: Venus-rGBD (O’Neill et al., 2018), mCherry 2xrGBD (Davenport et al., 2016), delCMV-EGFP-rGBD (Graessl et al., 2017) and 3xGFP-rGBD (Bement et al., 2015), but no comparison was published.

In order to understand and to expand the potential of relocation-based RhoA sensors, we first systematically compared and subsequently optimized several RhoA relocation sensors in cell-based assays. We quantified their relocation efficiency, checked their specificity for RhoA in comparison to Cdc42 and Rac1 and finally, we showcased their potential by visualizing endogenous RhoA activity in human endothelial cells.

## Results

### Optimizing the rGBD relocation RhoA sensor

To optimize the relocation RhoA biosensor we tested it in a cell-based assay. The eGFP-rGBD biosensor consists of an enhanced green fluorescent protein (eGFP) and a rhotekin G protein binding domain (rGBD). It reports active, i.e. GTP-bound, endogenous RhoA at the plasma membrane of *Xenopus* oocyte during wound healing (Benink & Bement, 2005). We verified the performance in HeLa cells, overexpressing the histamine 1 receptor (H1R), as we have previously demonstrated, activation of H1R by histamine activates RhoA in HeLa cells (Unen et al., 2016). In a resting HeLa cell, the rGBD sensor localizes in the cytosol. Upon histamine addition, it relocalizes to the membrane and it relocalizes to the cytosol when pyrilamine, a histamine antagonist, is added (**Figure 1A and 1C, Supplemental Movie 1**,**2**). However, the relocation of the original, single rGBD monomeric fluorescent protein sensor is hardly detectable. To optimize the rGBD sensors by increasing the avidity, we constructed single, double and triple rGBD mNeonGreen fusions. A triple mNeonGreen single rGBD version was created, to increase the brightness of a single sensor. A dimericTomato single and double rGBD sensor was generated to study the influence of a dimeric fluorescent protein on the sensor. We compared the change in cytosolic intensity of the sensor upon histamine addition, for different versions of the rGBD sensor (**Figure 1B**). We found that the change in cytosolic intensity increased with each added rGBD domain. The change from a single to a triple mNeonGreen did not change the performance of the sensor. Interestingly, a dimericTomato single rGBD sensor localizes as well as a monomeric fluorescent protein double rGBD sensor. So, a dimeric fluorescent protein provides another way to improve the performance of the location sensor by doubling the number of binding domains. We conclude that the dimericTomato-2xrGBD sensor shows the best relocation efficiency, with a median change in cytosolic intensity of close to 50%.

**Figure 1.**
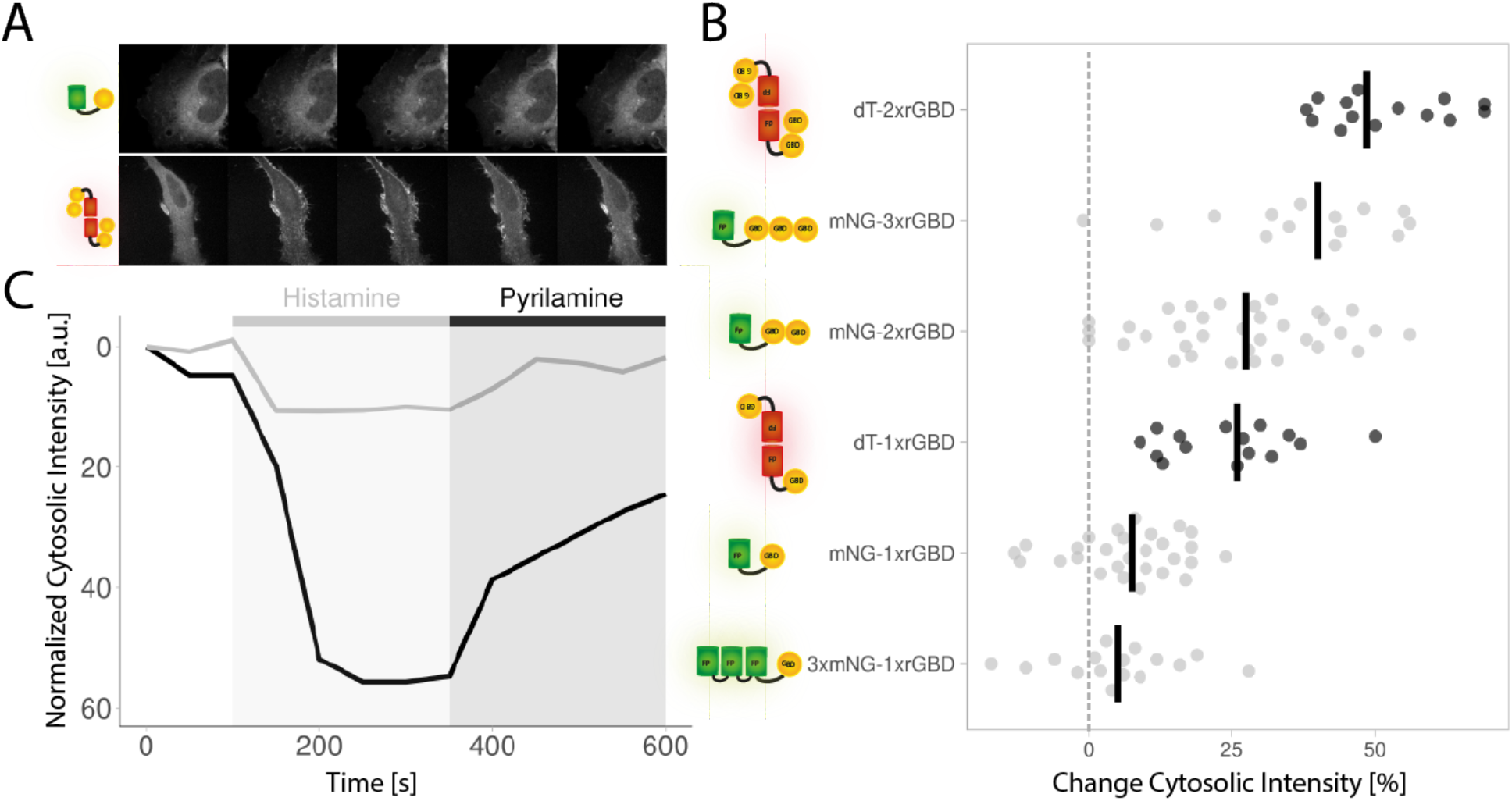
The optimized dimericTomato-2xrGBD RhoA sensor has the best location efficiency. **A**) Still images of a HeLa cells expressing the CMVdel-mNeonGreen-1xrGBD RhoA sensor (upper panel) or CMVdel-dimericTomato-2xrGBD RhoA sensor (lower panel) and H1R (not shown) which were stimulated with 100 μM histamine after 150 s and 10 μM pyrilamine after 350 s. **B**) Change in cytosolic intensity for mNeonGreen-1xrGBD/ - 2xrGBD/ -3xrGBD, 3xmNeonGreen-1xrGBD and dimericTomato-1xrGBD/ -2xrGBD in Hela cells expressing H1R, upon stimulation with 100 μM histamine. Each dot represents an individual cell. The median of the data is shown as vertical, black line and the grey dashed line indicates no change in cytosolic intensity. The number of samples per condition is: 3xmNG-1xrGBD=16, mNG-1xrGBD=28, dT-1xrGBD=15, mNG-2xrGBD=34, mNG-3xrGBD=14, dT-2xrGBD=14. **C**) Time traces of the normalized cytosolic intensity for the displayed cells for the mNeonGreen-1xrGBD sensor in grey and for the dimericTomato-2xrGBD sensor in black. Abbreviations: mNG: mNeonGreen, dT: dimericTomato, rGBD: rhotekin G protein binding domain

### Coexpression comparison of RhoA location sensors

We then performed a co-expression experiment, to directly compare the relocalization of the dimericTomato-2xrGBD sensor to the mTurquoise2-1xrGBD sensor in the same HeLa cell (**Figure 2A, Supplemental Movie 3**), avoiding cell-to-cell heterogeneity. Where the single rGBD sensor only shows a 10% drop of cytosolic intensity compared to baseline, the optimized dimericTomato-2xrGBD sensor shows a 40% decrease in intensity (**Figure 2B**). We also compared the dimericTomato-2xrGBD sensor to an alternative localization-based sensor for RhoA (**Figure 2C, Supplemental Movie 4**), which utilizes the AHD and PH domain of anillin (Munjal et al., 2015; Piekny & Glotzer, 2000). The anillin sensor AHD+PH showed a 15% decrease in cytosolic intensity (**Figure 2D**), but it also relocalizes to striking punctuate structures upon histamine stimulation. These structures did not seem to represent local, high activity of RhoA, as the optimized rGBD sensor in the same cell showed no such locally clustered RhoA activation, but rather a homogenous activation at the membrane and a 60% drop in cytosolic intensity. Similar punctuate structures were observed in endothelial cells, when stimulated with the strong RhoA activator thrombin (**Supplemental Movie 5**). Concluding, the optimized dimericTomato-2xrGBD sensor outperforms two existing RhoA relocation sensors in a direct comparison.

**Figure 2.**
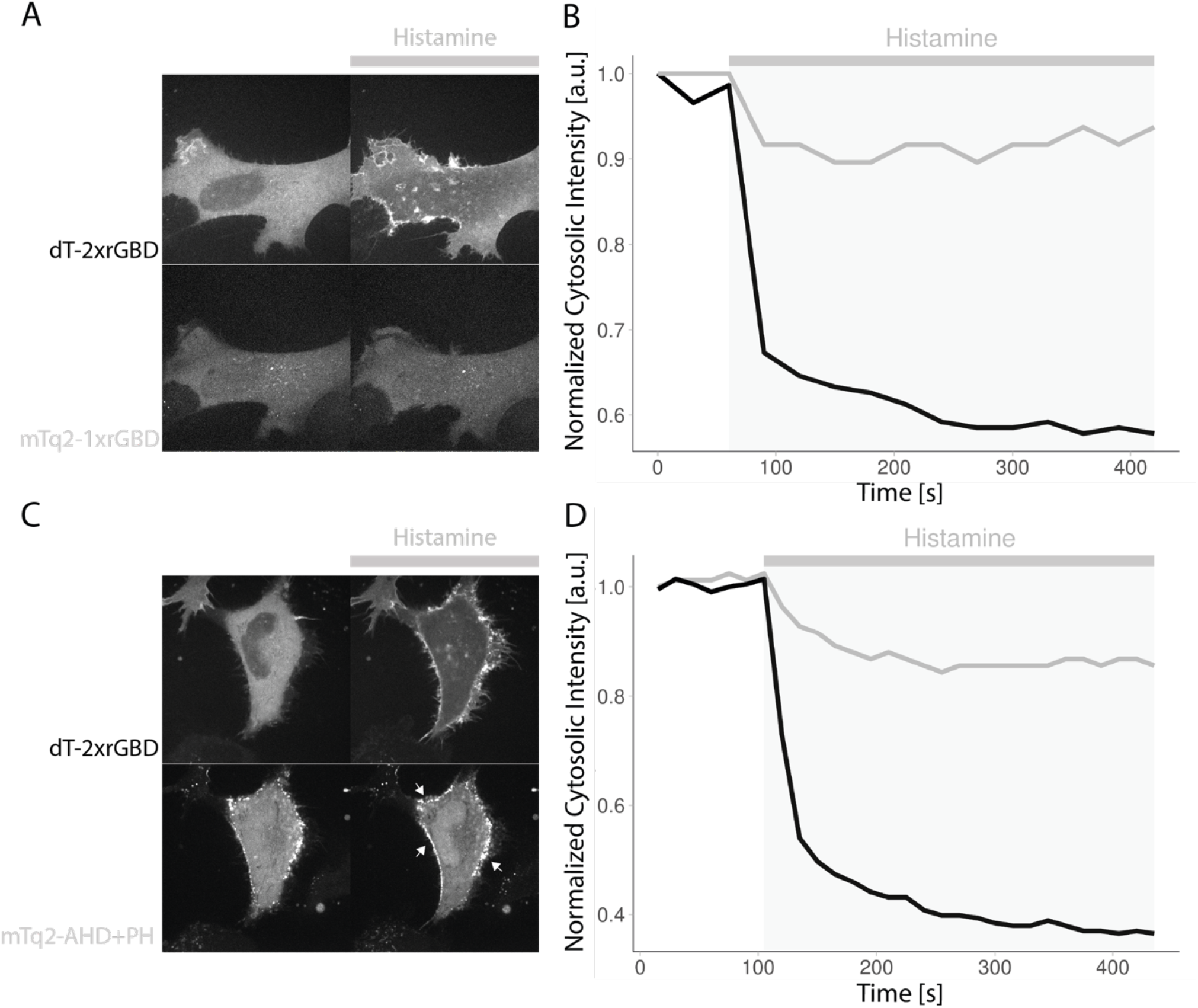
The optimized dimericTomato-2xrGBD RhoA sensor localizes more efficiently than the published single rGBD RhoA sensor and anillin RhoA sensor. **A)** Still images of a HeLa cell expressing H1R (not shown) and CMVdel-dimericTomato-2xrGBD (upper panel) and CMVdel-mTurquoise2-1xrGBD (lower panel) before (left) and 5 min after (right) stimulation with 100 μM histamine. **B)** Traces of the normalized cytosolic intensity for CMVdel-dimericTomato-2xrGBD (black) and CMVdel-mTurquoise2-1xrGBD (grey) for the displayed cell. **C**) Still images of a HeLa cell expressing H1R (not shown) and CMVdel-dimericTomato-2xrGBD (upper panel) and CVM-mTurquoise2-Anillin(AHD+PH) (lower panel) before (left) and 5 min after (right) stimulation with 100 μM histamine. **D**) Traces of the normalized cytosolic intensity for CMVdel-dimericTomato-2xrGBD (black) and CMV-mTurquoise2-Anillin(AHD+PH) (grey) for the displayed cell.

### Specificity of rGBD for RhoA

Next, we wanted to examine the selectivity of the sensor for RhoA in comparison to other Rho GTPases in living cells. To this end, we generated nuclear localized, constitutively active Rho GTPases (H2A-mTurquoise2-RhoAG14V-ΔCaaX, H2A-mTurquoise2-Rac1G12V-ΔCaaX and H2A-mTurquoise2-Cdc42G12V-ΔCaaX), a strategy that was used before (Bery et al., 2019). The H2A histone tag, in combination with the removal of the CaaX box, allows the construct to completely localize in the nucleus, otherwise it is partly directed to the plasma membrane. With this approach, binding affinity can be assessed by colocalization of the location-based sensor with the applicable Rho GTPase. We co-expressed these constitutively active Rho GTPases in Hela cells with CMVdel-dimericTomato-2xrGBD or CMVdel-mScarlet-I-1xrGBD and measured the intensity of the sensor in the nucleus in comparison to the cytosol (**Figure 3A and 3B**). The rGBD sensors solely colocalized in the nucleus with RhoA but not with Rac1 and Cdc42, indicating that rGBD specifically binds constitutively active RhoA. This is in line with previous studies in cell extracts and bacterial lysate (Reid et al., 1996; Ren et al., 1999). Comparing the original single rGBD sensor with the dimericTomato-2xrGBD sensor in colocalization with RhoA G14V, a higher nuclear to cytosolic intensity ratio for the multi-domain sensor was detected.

**Figure 3.**
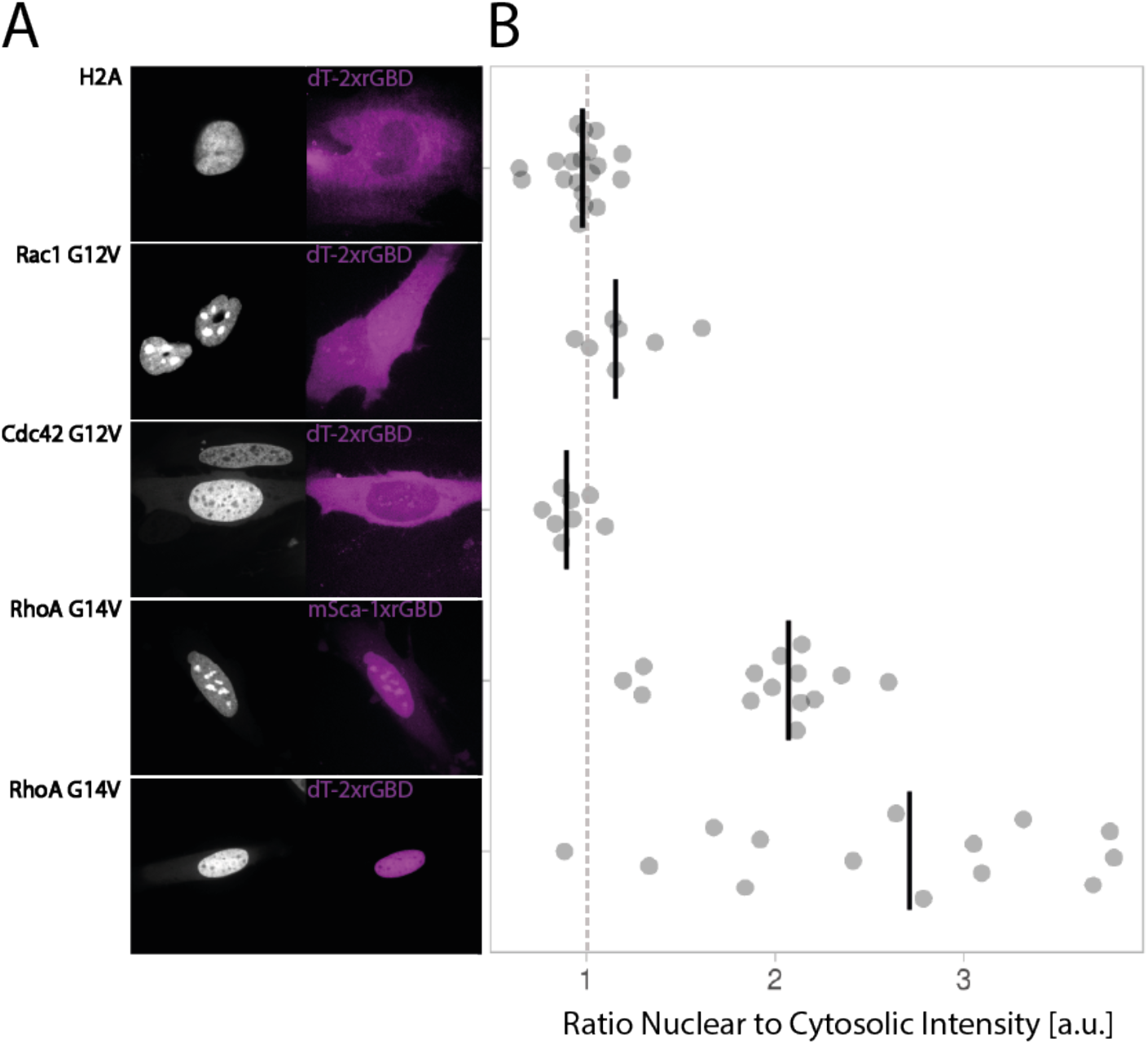
The DimericTomato-2xrGBD RhoA sensor is specific for RhoA. **A**) Colocalization of dimericTomato-2xrGBD or mScralet-I-1xrGBD with H2A-mTurquoise2-Rho GTPase-ΔCaaX in HeLa cells, for control H2A-mTurquoise2, Rac1, Cdc42, and RhoA. **B**) Rho GTPase specificity, represented by the ratio of dimericTomato-2xrGBD or mScralet-I-1xrGBD intensity in the nucleus to cytosol in H2A-mTurquoise2-Rho GTPase-ΔCaaX expressing HeLa cells. Each dot represents an individual cell. The dashed line indicates a ratio of one. The median of the data is shown as a vertical, black line. The number of samples per condition is: RhoA-dT2xrGBD=14, RhoA-mSca-1xrGBD=14, Cdc42-dT2xrGBD=8, Rac1-dT-2rGBD=7, H2A-dT2xrGBD=19.

### Attempt to utilize G protein binding domains of Anillin and PKN1 to create a RhoA location sensor

Given the successful improvement of the rGBD-based biosensor by increasing the number of binding domains, we explored whether the same strategy can be applied to the G protein binding domains from PKN1 and Anillin. The G protein binding domain of PKN1 (pGBD), used in DORA RhoA FRET sensor (Van Unen et al., 2015), was used as starting material for a relocation sensor. Moreover, a published relocation sensor based on Anillin contains, next to a G protein binding domain, also a C2 and a PH domain and localizes in punctuate structures which do not represent RhoA activity (**Figure 2C**,**Supplemental Movie 4 and 5**) (Munjal et al., 2015; Piekny & Glotzer, 2000). Here, we used only the G protein binding domain of Anillin (aGBD) as a basis for another sensor.

Using the same strategy as for the rGBD sensors, single, tandem and triple and dimericTomato versions of the sensor were created (CMVdel-mNeonGreen-1xpGBD/ -2xpGBD/ -3xpGBD and CMVdel-dimericTomato-2xpGBD) and tested in H1R-expressing Hela cells by stimulating endogenous RhoA with histamine. None of the pGBD sensors showed a clear membrane localization upon stimulation with histamine (**Figure 4A**). The increase in cytosolic intensity observed in some cells, seems to be caused by changes in cell shape. Nevertheless, when CMVdel-dimericTomato-2xpGBD is co-expressed with H2A-mTurquoise2-RhoAG14V-ΔCaaX (constitutively active and nuclear located RhoA) in Hela cells, pGBD accumulated in the nucleus (**Figure 4B**), indicating that pGBD does bind constitutively active RhoA.

**Figure 4.**
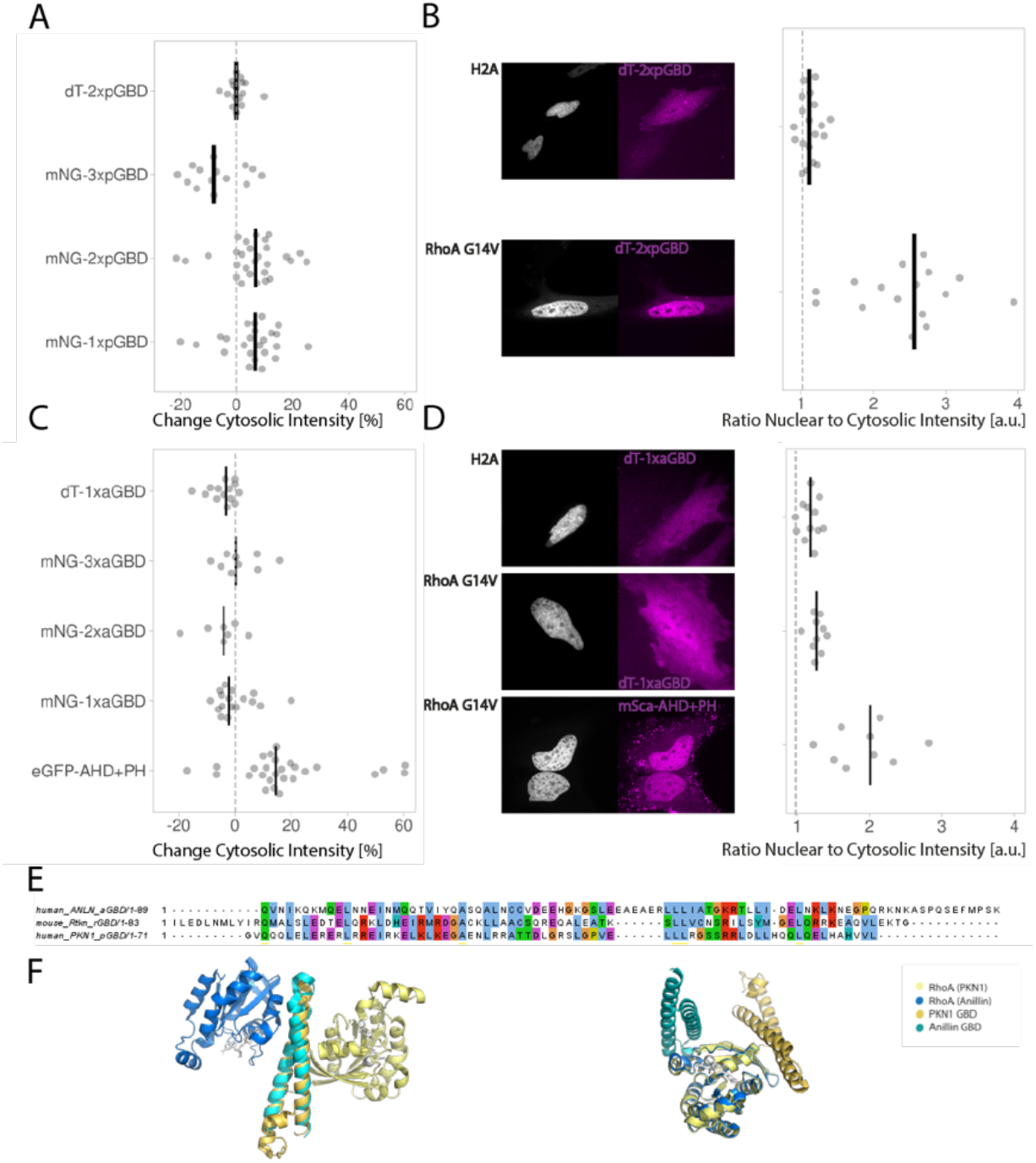
G Protein Binding Domains of Anillin and PKN1 are not suitable for a relocation RhoA sensor. **A**) Change in cytosolic intensity for CMVdel-mNeonGreen-1xpGBD/ -2xpGBD/ -3xpGBD, CMVdel-dimericTomato-2xpGBD coexpressed with H1R in HeLa cells upon stimulation with 100 μM histamine. The dashed line represents no change in cytosolic intensity. Each dot represents an individual cell. The median of the data is shown as vertical, black line. The number of samples per condition is: dT-2xpGBD=16, mNG-1xpGBD=24, mNG-2xpGBD=28, mNG-3xpGBD=14. **B**) Colocalization of H2A-mTurquoise2-RhoAG14V-ΔCaaX or control H2A-mTurquoise2 with dimericTomato-2xpGBD in HeLa cells. RhoA binding, represented by the ratio of sensor intensity in the nucleus to cytosol in H2A-mTurquoise2-RhoAG14V-ΔCaaX expressing HeLa cells. The dashed line indicates a ratio of one. Each dot represents an individual cell. The median of the data is shown as a vertical, black line. The number of samples per condition is:H2A-dT-2xpGBD=19, RhoA-dT2xpGBD=16. **C**) Change in cytosolic intensity for CMVdel-mNeonGreen-1xaGBD/ -2xaGBD/ -3xaGBD, CMVdel-dimericTomato-1xaGBD coexpressed with H1R in HeLa cells upon stimulation with 100 μM histamine. The dashed line represents no change in cytosolic intensity. Each dot represents an individual cell. The median of the data is shown as vertical, black line. The number of samples per condition is: dT-1xaGBD=14, eGFP-AHD+PH=27, mNG-1xaGBD=17, mNG-2xaGBD=7, mNG-3xaGBD=9. **D**) Colocalization of H2A-mTurquoise2-RhoAG14V-ΔCaaX or control H2A-mTurquoise2 with dimericTomato-1xaGBD and mScarlet-I-AHD+PH in HeLa cells. RhoA binding, represented by the ratio of sensor intensity in the nucleus to cytosol in H2A-mTurquoise2-RhoAG14V-ΔCaaX expressing HeLa cells. The dashed line indicates a ratio of one. The median of the data is shown as a vertical, black line. The number of samples per condition is: H2A-aGBD=12, RhoA-aGBD=10, RhoA-AHD=9. **E**) Amino acid sequence alignment for aGBD, rGBD and pGBD by MUSCLE depicted with the clustalX color code. **F**) On the left a structural alignment of PKN1 and Anillin by their RhoA binding domains. On the right a structural alignment of PKN1 and anillin by RhoA, showing the two binding positions at the RhoA molecule. Anillin and the bound RhoA are depicted in dark and light yellow, respectively. PKN1 and the bound RhoA are depicted in light and dark blue, respectively. (PDB: Anillin = 4xOI, PKN1 = 1cxz).

For the original Anillin AH+PH sensor, a variety of responses can be observed when H1R expressing HeLa cells are stimulated with histamine. A small pool of cells showed a cytosolic intensity change between 50% and 60 %. A large pool showed a cytosolic intensity change of around 15%. Plus, the sensor localized in a clustered, non-homogenous manner in all these cells. However, for the aGBD-based sensors (CMVdel-mNeonGreen-1xaGBD/ -2xaGBD/ -3xaGBD and CMVdel-dimericTomato-1xaGBD), no localization to the membrane upon histamine stimulation was observed (**Figure 4C**). Additionally, when CMVdel-dimericTomato-1xaGBD and mScarlet-I-AHD+PH are coexpressed with H2A-mTurquoise2-RhoAG14V-ΔCaaX (constitutively active and nuclear located RhoA) in Hela cells only the AHD+PH (aGBD+C2+PH) construct localizes with the active RhoA (**Figure 4D**). The aGBD by itself did not localize with the RhoA in the nucleus, indicating that it is not able to bind RhoA without the C2 and PH domain.

Given the different behavior of aGBD, pGBD and rGBD, we examined their amino acid sequence and structure. The amino acid alignment of the GBDs showed conserved hydrophobic residues (**Figure 4E**). It also showed that all three domains contain the leucine repeats, which have been shown to interact with RhoA at the example of pGBD (Maesaki et al., 1999). The superimposed crystal structures of aGBD and pGBD binding to RhoA-GTP showed a good overlap of the GBDs and revealed two different binding sites for RhoA (**Figure 4F**). These sites have previously been described for pGBD (Maesaki et al., 1999) and this may be a general feature of GBDs. Unfortunately, no crystal structure is available for rGBD. The amino acid sequence and structure did not provide a clear explanation for the different behavior of the three GBDs.

In conclusion, the attempt to create a RhoA location sensor from Anillin and PKN1 was not successful.

### Visualizing endogenous RhoA at the Golgi

In all of the previous experiments, endogenous RhoA activity was detected at the plasma membrane. A subset of RhoA is known to localize at the Golgi apparatus membrane (Zilberman et al., 2011). We challenged the sensitivity of the optimized RhoA biosensor to detect activity of this smaller RhoA fraction. To examine this, we used a rapamycin-induced hetero dimerization system to recruit the dbl homology (DH) domain, of the RhoA activating GEF p63, to the membrane of the Golgi apparatus. We chose the plasma membrane as a positive control and mitochondria as negative control. Recruiting p63-DH to the plasma membrane caused a clear increase in RhoA biosensor intensity at the plasma membrane (**Figure 5, Supplemental Movie 6**). Recruiting p63-DH to the Golgi apparatus is followed by a slight increase of RhoA biosensor intensity at this organelle (**Figure 5, Supplemental Movie 7**). Recruiting p63-DH to mitochondria did not result in a clear increase of the RhoA biosensor intensity (**Figure 5, Supplemental Movie 8**). In summary, the dimericTomato-2xrGBD sensor is able to visualize activity of the small RhoA fraction at the Golgi apparatus.

**Figure 5.**
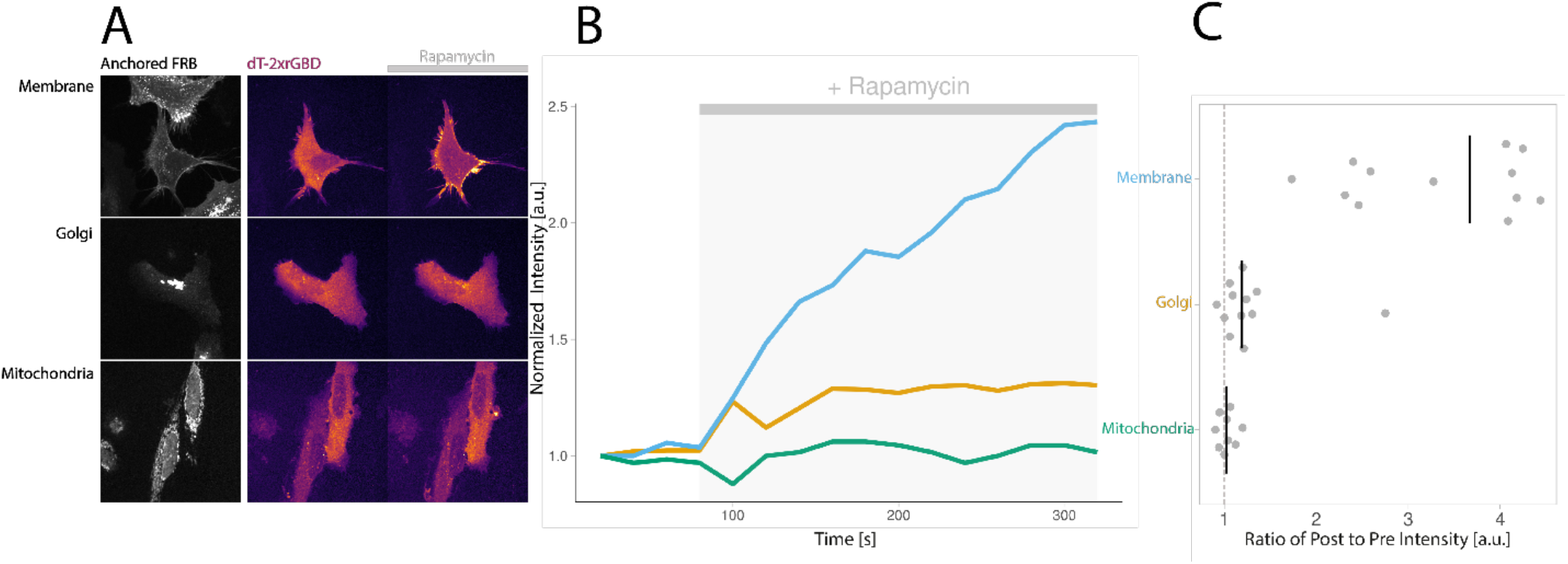
The optimized dimericTomato-2xrGBD sensor shows organelle specific RhoA activity. **A**) Still images of HeLa cells expressing FRB anchored to the membrane, Golgi and mitochondria (first column), FKBP-p63-DH, localization of the dimericTomato-2xrGBD sensor pre activation (second column) and post activation with 100 nM rapamycin (third column). **B**) Time traces of the dimericTomato-2xrGBD sensor intensity for the displayed cells at the indicated location. **C**) Localization of the dimericTomato-2xrGBD sensor represented by the ratio of post to pre rapamycin activation intensity. The median is indicated by a vertical, black line and the ratio of 1 representing no accumulation of the sensor is indicated by a dashed grey line. The number of samples per condition is: Golgi=12, Membrane=12, Mitochondria=9.

### Visualizing endogenous RhoA activity

Finally, we used the optimized biosensor to visualize endogenous RhoA activation under physiological conditions, with attention to subcellular relocalization in combination with RhoA mediated processes. Therefore, we studied a HeLa cell expressing the dimericTomato-2xrGBD RhoA biosensor going through cell division (**Figure 6A, Supplemental Movie 9**). The sensor localized clearly at the cleavage furrow where accumulation of active RhoA has been reported (Piekny & Glotzer, 2000). Next, we examined human endothelial cells expressing the dimericTomato-2xrGBD RhoA biosensor and observed random migration, where localization of the sensor at the cell edge is followed by contraction of that cell edge (**Figure 6B, Supplemental Movie 10**). Both of these results confirmed that the RhoA biosensor relocalizes to cellular structures where RhoA activity is expected.

**Figure 6.**
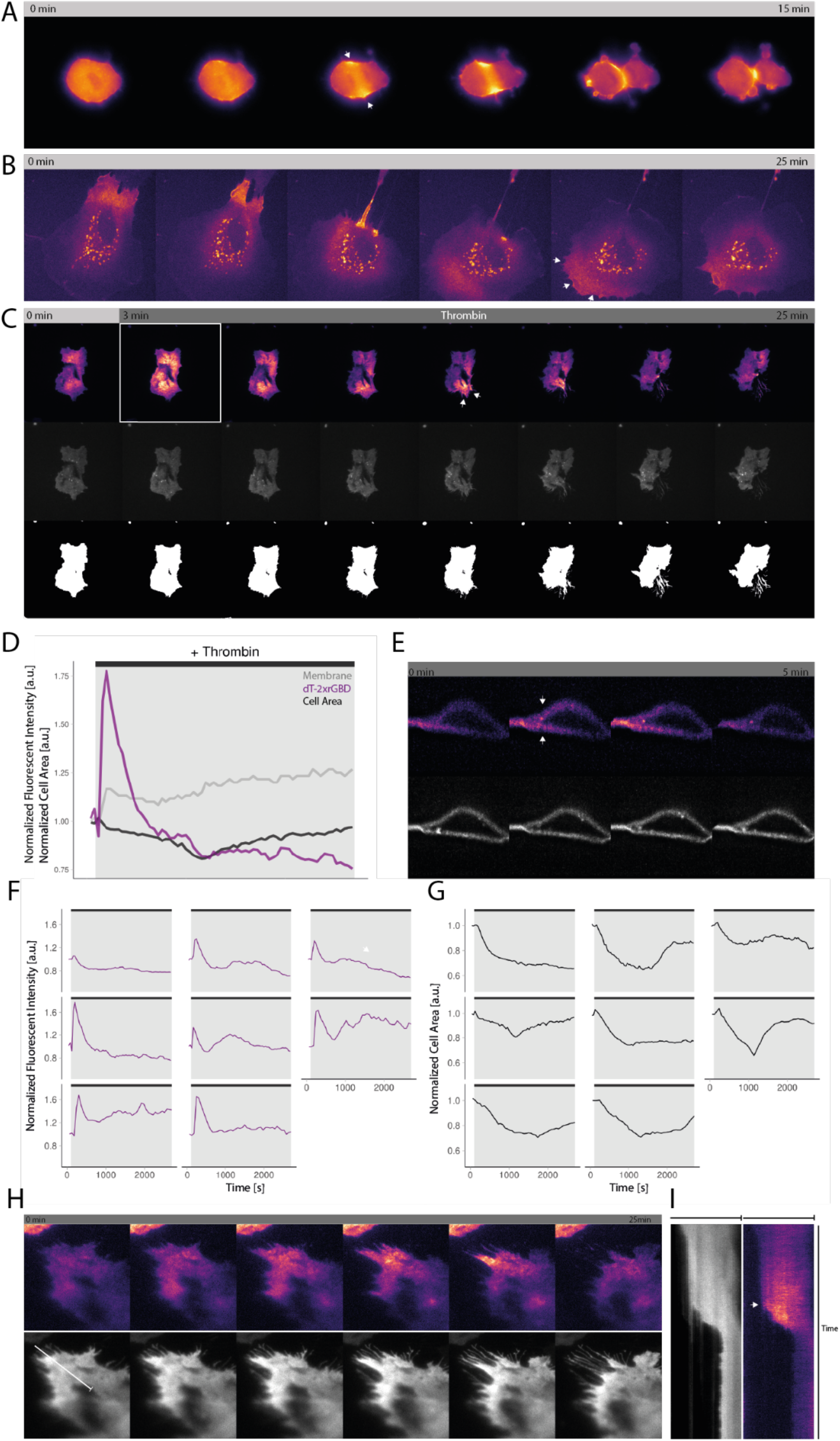
Visualization of endogenous RhoA activity. **A**) Wide field images of a Hela cell expressing dimericTomato-2xrGBD going through cell division. Arrows indicate intensity increase at the location that will become the cleavage furrow. **B**) Spinning disk images of a BOEC transiently expressing dimericTomato-2xrGBD randomly migrating after cell division. Arrows indicate contracting cell edge with increased intensity. **C**) TIRF images of a cbBOEC stably expressing dimericTomato-2xrGBD (upper panel) and mTq2-CaaX (middle panel) stimulated with 1 U/ml human α-thrombin. The lower panel shows the cell area as a binary image. The white frame indicates the moment of global dimericTomato-2xrGBD intensity increase. Arrows indicate local dimericTomato-2xrGBD intensity increase followed by local contraction. **D**) Time trace of the normalized fluorescent intensity for the dimericTomato-2xrGBD sensor (purple) and the mTq2-CaaX membrane label (grey) for the cell depicted in C. The normalized cell area (black) was measured for the white region shown in the lower panel of C. The same region was used as ROI for the fluorescent intensity measurements. **E**) Lattice light sheet cross section images of a HUVEC stably expression dimericTomato-2xrGBD (upper panel) and mTq2-CaaX (lower panel). The cell was stimulated with 2.5 µM nocodazole 10 min prior to the imaging. Arrows indicate dimericTomato-2xrGBD intensity increase at basal and apical plasma membrane. **F**) Time traces of the normalized fluorescent intensity of dimericTomato-2xrGBD stably expressed in single BOECs, imaged with TIRF microscopy, stimulated with 1 U/ml human α-thrombin after 120 s. The stimulation is indicated by the black bar and the grey box. **G**) Time trace of the normalized cell area of BOECs. The measurements correspond to the intensity measurements in H the same position of the graph represents the same cell. **H**) TRIF images of a cbBOEC stably expressing dimericTomato-2xrGBD (upper panel) and mTq2-CaaX (lower panel). The cell was stimulated with 1 U/ml human α-thrombin 5 min prior to the imaging. Arrows indicates dimericTomato-2xrGBD intensity increase followed by contraction of the cell edge. **I**) Kymograph of the cell depicted in F along the white line. The fluorescent intensity of mTq2-CaaX is depicted in grey and the fluorescent intensity of dimericTomato-2xrGBD is depicted with the LUT mpl-magma. Arrows indicates dimericTomato-2xrGBD intensity increase followed by contraction.

Then we turned to TIRF microscopy to specifically image the dimericTomato-2xrGBD at the basolateral membrane of an endothelial cell. We co-expressed a membrane marker, mTurquoise2-CaaX, to correct for intensity changes unrelated to sensor relocation. Endothelial cells were stimulated with human α-thrombin to activate endogenous receptors (**Figure 6C, Supplemental Movie 11**). We measured a peak in RhoA biosensor intensity of roughly 75% increase globally over the whole basolateral plasma membrane, within seconds after stimulation with human α-thrombin (**Figure 6D**). The sensor intensity decreased to base level after approximately 6 min, this is in line with previous research (Heemskerk et al., 2016). The cell area decreased approximately 20% reaching the minimum at 20 min after stimulation, followed by recovering of the cell area to the original size over the course of 40 min after stimulation, which is in line with what is known about thrombin-responsive endothelial cells (Timmerman et al., 2015). The membrane marker showed a relatively small increase in intensity after stimulation and the curve did not show the same pattern as the RhoA biosensor intensity curve. Therefore, we conclude that the increase in RhoA biosensor intensity is caused by relocalization. The global increase in RhoA biosensor intensity followed by cell contraction was robustly observed for multiple cells (**Figure 6F and 6G**).

Since TIRF microscopy only visualizes RhoA activity at the basolateral membrane, we imaged a cross section of the cell using lattice light sheet microscopy (**Figure 6E, Supplemental Movie 12 and 13)**. These cross sections showed the increase in RhoA biosensor intensity at the basolateral as well as at the apical plasma membrane. The (control) membrane marker showed no visible increase.

Following the first global increase in RhoA biosensor intensity upon human α-thrombin stimulation, we also observed local increase followed by retraction in this cell area (**Figure 6H, Supplemental Movie 14**). Kymograph analysis showed, the increase of the RhoA biosensor intensity is followed by a retraction of the cell’s periphery, and the increase in intensity is not reflected in the membrane marker (**Figure 6I)**. The local cell contraction confirmed that the RhoA biosensor indeed sensed endogenous active RhoA with high spatial resolution. Cell contraction is the expected cellular response upon RhoA activation (Ridley & Hall, 1992).

It is important to note that even though the sensor binds endogenous RhoA, it allows for sensitive cellular responses such as full cell division and cell contraction. Moreover, our data revealed that the RhoA biosensor displays RhoA activity at subcellular locations where RhoA activity is expected, and appears mostly independent of fluorescent intensity measured by a separate membrane marker.

## Discussion

RhoA relocation sensors have been used for more than a decade, but their use has been mostly limited to *Xenopus* oocytes, macrophages and *Drosophila* embryos (Benink & Bement, 2005; Jiang & Harris, 2019; Mason et al., 2016; O’Neill et al., 2018). Using the original eGFP-rGBD sensor in HeLa cells, we observed only a subtle relocalization. The reason for this poor performance in mammalian cell cultures has been unclear. We systematically increased the avidity by increasing the number of binding domains. This resulted in a drastic increase in the relocalization efficiency. Moreover, we demonstrated that the use of a dimerizing fluorescent protein is an efficient strategy to increase the avidity. The improved dimericTomato-2xrGBD sensor relocalizes with higher efficiency than the rGBD and AHD+PH sensors in a direct comparison. Consequently, our data show that it is the preferred relocation sensor to study endogenous RhoA activity in mammalian cells. We show that the dimericTomato-2xrGBD sensor can visualize endogenous RhoA activity with high spatial resolution, in expected locations such as the cleavage furrow of a dividing HeLa cell and prior and during cell retraction in endothelial cells.

The systematic optimization of the rGBD probe by increasing the avidity was successful. This strategy, to utilize multiple repeating domains has also been effective for a PH domain based lipid sensor (Goulden et al., 2018). The avidity of this lipid sensor was increased which each added PH domain, indicating a cooperative effect, which decreases the off-rate of binding the plasma membrane. However, increasing the number of binding domains to generate relocation sensors from other RhoA binding domains, i.e. aGBD and pGBD, was not successful. The previously reported AHD+PH RhoA sensor was derived from anillin, a scaffold protein that connects RhoA, actin and myosin, especially during cell division (Piekny & Glotzer, 2000). When we utilized only the aGBD domain and fused it to a fluorescent protein, we obtained a probe that does not colocalize with active RhoA, whereas AHD+PH does. This result is in line with the notion that only the synergistic action of aGBD, C2 and PH enables anillin to bind active RhoA and to localize at the plasma membrane (Sun et al., 2015). However, we noticed that the AHD+PH sensor, containing aGBD, C2 and PH domain, localizes in a punctate manner. These ‘dots’ were observed in both HeLa cells and endothelial cells and were only observed with the AHD+PH RhoA sensor. As aGBD does not localize in puncta, it seems that the localization is caused by domains other than of the RhoA binding domain, i.e. the C2- and/or PH-domain.

Strikingly, the pGBD that works in the DORA RhoA FRET-based sensor, does not work as a relocation sensor. The DORA FRET-based sensor for RhoA uses the pGBD domain from PKN1, a member of the protein kinase C-related family of serine/ threonine protein kinases, mediating a Rho GTPase-dependent signaling pathway (Lim et al., 2006). Our data show that the pGBD location sensor is able to bind active RhoA in a living cell assay. However, it does not relocalize upon RhoA activation and the addition of multiple bindings domains or a dimeric fluorescent protein does not improve its avidity.

Looking at the amino acid sequence of aGBD, pGBD, rGBD and their structures did not give an explanation why rGBD works in a relocation sensor and the other two do not. The function of rhotekin is not clear, it seems to lock RhoA in the GTP bound state (Ito et al., 2018; Reid et al., 1996). We can only speculate that rhotekin has a stronger binding affinity for active RhoA than anillin and PKN1 have. This idea is supported by a mass spectrometry-based study for active RhoA binders, where rhotekin scores higher than anillin and PKN1 (Gillingham et al., 2019). This difference in affinity might explain its ability to function as a relocation sensor for RhoA. The unimolecular FRET-based sensors that consist of RhoA and a GBD probably require a lower binding affinity. This may explain why pGBD can be used in a FRET-based sensor but not as a relocation sensor. Unfortunately, we have not found a ‘recipe’ that would allow us to convert any RhoA binding protein into a relocation sensor.

Comparing relocation sensors to FRET sensors, both have their own advantages and disadvantages. Relocation sensor data lack the semi quantitative aspect of ratio FRET measurements. Moreover, one has to consider that morphological changes can lead to fluorescent intensity changes (Dewitt et al., 2009). The ratio metric approach of the FRET sensor accounts for these morphological intensity changes, provided the intensity change is equal in both channels. A solution for intensity changes unrelated to the relocation of the biosensor is to co-image an inert plasma membrane bound fluorescent marker. The optimized RhoA biosensor intensity increase was mostly unrelated to membrane fluorescent intensity increase. The specificity of the relocation sensor is determined by the binding specificity of the GBD. The rGBD binds the three homologues RhoA, B and C but not to Rac1 and Cdc42 (Ren et al., 1999). To further improve the binding specificity, one either needs to screen for specific RhoA binders or design an artificial GBD. The latter has been done in form of an anti RhoA-GTP nanobody (Keller et al., 2019).

Furthermore, the relocation sensor requires confocal microscopy or TIRF microcopy to spatially separate the bound from unbound probe, whereas FRET measurements are usually performed with widefield microscopes. However, the former mentioned techniques usually offer the higher resolution. Here we presented previously unachieved visualization of RhoA activity at subcellular resolution. We observed local activation of RhoA at the Golgi which was not possible with the DORA RhoA FRET sensor (Van Unen et al., 2015), indicating a higher sensitivity of the relocation sensor. It is worth noting that the operating principal of the two sensor types is different. Whereas the RhoA FRET sensor is a read out of endogenous GEF activity, the RhoA location sensor detects endogenous RhoA activity directly. The optimized RhoA relocation sensor visualized endogenous RhoA activity at its true location in the cell with great spatial detail.

Visualizing the endogenous RhoA activity may interfere with the biological role of RhoA, as the sensor binds endogenous RhoA and may compete with natural effectors of RhoA. As an example, the rGBD has been used as RhoA inhibitor in zebrafish (Yoo et al., 2010). To limit the perturbation, the sensor should be expressed at a low level to allow RhoA signaling. We demonstrate that low expression of the biosensor, through the truncated CMV promotor, did not inhibit cell division and cell edge retraction. Plus, endothelial cells expressing the sensor still show the typical reaction of contracting followed by spreading, when stimulated with thrombin. Low expression results in a low fluorescent signal of the sensor. The dimericTomato-2xrGBD sensor has the advantage, that when it forms a dimer, one sensor unit contains two dimerTomato molecules and 4 rGBDs. To enhance the brightness per sensor molecule, one could introduce a triple fluorescent protein in combination with multiple rGBD, rather than searching for a strongly dimeric fluorescent protein. Using a triple fluorescent protein gives the choice of any characterized fluorescent protein which will be an advantage for multiplexing. If cells express RhoA at a higher level or if their shape allows to see a thicker part of the membrane in the Z plane, the relocation sensor will localize more clearly, which might explain the different performance in different cellular systems. Another way to circumvent the influence of the cell shape on the location efficiency is TIRF microscopy. This has been done for the rGBD sensor by two groups in U-2 OS cells and in *Drosophila* Schneider (S2) cells (Graessl et al., 2017; Verma & Maresca, 2019). While we were able to image dimericTomato-2xrGBD relocalization with a spinning disk set up, TIRF microscopy provides a better signal to noise ratio and less bleaching. These properties can be of great advantage while working with low expression of the biosensor.

The dimericTomato-2xrGBD genetically encoded, single color fluorescence biosensor gives the opportunity to measure endogenous RhoA activity with high spatial resolution. Single color relocation sensors are ideal candidates for multiplexing experiments. Plus, the growing field of optogenetics is in need of single color biosensors to detect the effect of optogenetic perturbations. The conventional CFP-YFP FRET sensor is incompatible with most, blue light induced optogenetic tools. Another research group has shown that relocation sensors combine well with optogenetics, here for example, recruiting the RhoA GEF LARG with the iLID system to the membrane driving cell migration, RhoA activity measured with Venus rGBD (O’Neill et al., 2018).

To conclude, we succeeded in visualizing endogenous RhoA with high spatial and temporal resolution in living mammalian cells with the improved, single color rGBD RhoA relocation biosensor. We expect that the new probe will be a versatile tool to measure RhoA activity in living cells and tissue. Beyond, we imagine that multiplexing Rho GTPase relocation sensors will be key to understand complex cellular processes.

## Material and Methods

### Plasmid Construction

#### rGBD

GFP-rGBD was a gift from William Bement (Addgene plasmid # 26732). Reduced Expression GFP beta actin was a gift from Rick Horwitz & Tim Mitchison (Addgene plasmid # 31502). In this plasmid the base pairs 91-544 of the enhancer region in the CMV promoter are deleted, in the following the promoter will be called CMVdel. The rGBD was cut with BsrGI and XbaI and cloned into a demethylated and likewise digested mCherry-C1 vector. mCherry was replaced with mNeonGreen/ 3xmNeonGreen using the AgeI and BsrGI restriction sites. The insert mNeonGreen-rGBD and the backbone Reduced Expression GFP beta actin were digested with AgeI and MluI. The digested products were ligated to create CMVdel-mNeonGreen-1xrGBD. A tandem and a triple rGBD were created by PCR amplification of rGBD from CMVdel-mNeonGreen with primers shown in Table 1 and digestions by BsrGI, SalI. The backbone CMVdel-mNeonGreen1xrGBD, or CMVdel-mNeonGreen-2xrGBD respectively, were digested with BsrGI, AvaI. Backbone and insert were ligated to generate CMVdel-mNeonGreen-2xrGBD/-3xrGBD.

**Table 1.**
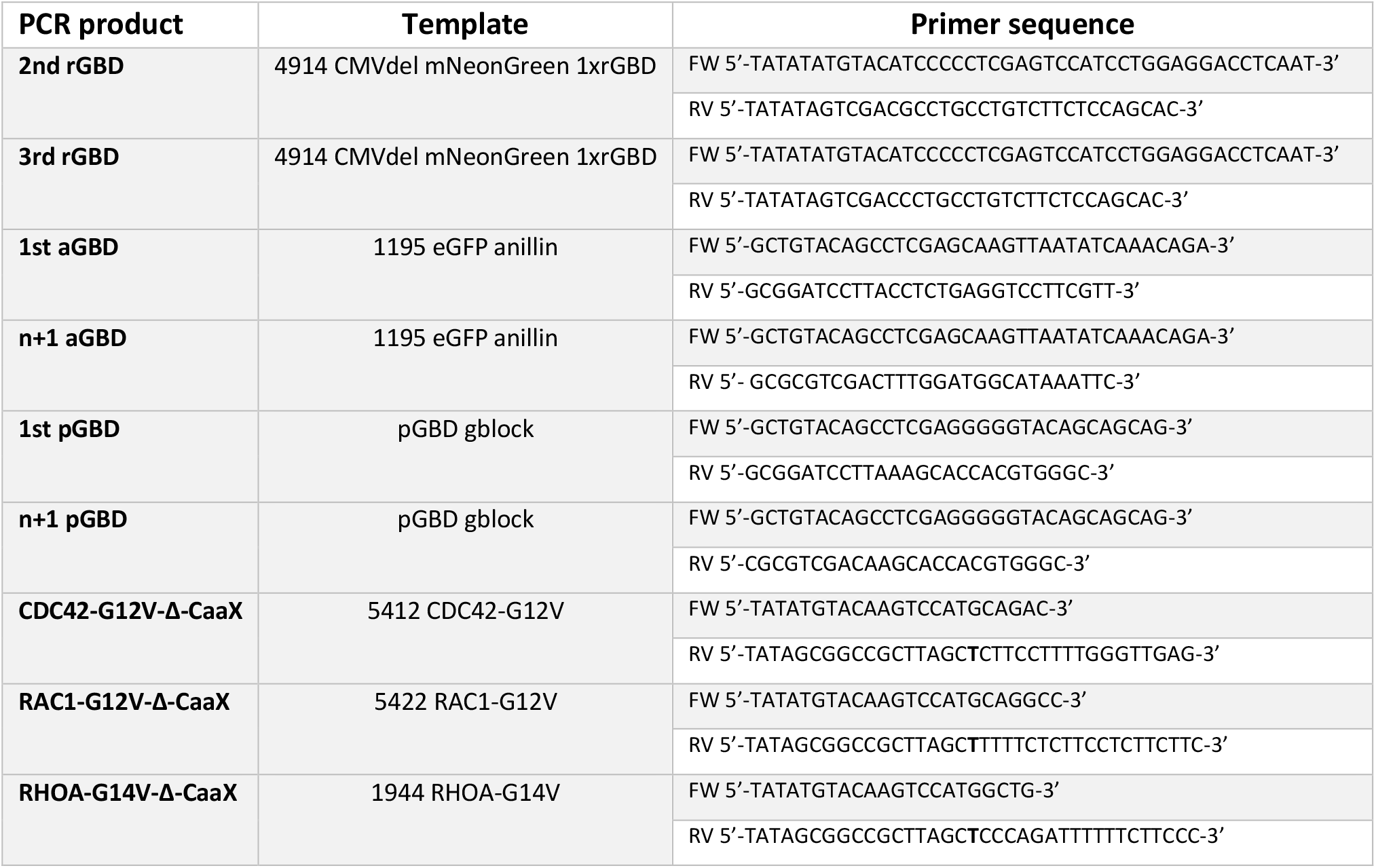
PCR primers for insert amplification.

**Table 2.** Plasmid names and cloning strategy. The sequences of the plasmid maps are available as supplementary information. The 4-digit number in the first column matches the filename of the plasmid sequence.

|  | NAME | BACK-BONE | RESTRICTION ENZYMES<br>BACKBONE | INSERT | RESTRICTION<br>ENZYMES INSERT |
| --- | --- | --- | --- | --- | --- |
| 4914 | CMVdel-mNeonGreen-1xrGBD | 4900 | AgeI, BsrGI | 4907 | AgeI, BsrGI |
| 5196 | CMVdel-dTomato-1xrGBD | 4914 | AgeI, BsrGI | 4595 | AgeI, BsrGI |
| 5215 | CMVdel-mScarlet-I-1xrGBD | 4914 | AgeI, BsrGI | 4604 | AgeI, BsrGI |
| 5214 | CMVdel-mTurquoise2-1xrGBD | 4914 | AgeI, BsrGI | 3703 | AgeI, BsrGI |
| 1199 | CMV-3xmNeonGreen-1xrGBD | 3877 | AgeI, BsrGI | 2376 | AgeI, BsrGI |
| 5380 | CMVdel-dTomato-2xrGBD | 4915 | AgeI, BsrGI | 4595 | AgeI, BsrGI |
| 4915 | CMVdel-mNeonGreen-2xrGBD | 4914 | BsrGI, Aval | 2nd rGBD | BsrGI, Sall |
| 4917 | CMVdel-mNeonGreen-3xrGBD | 4915 | BsrGI, Aval | 3rd rGBD | BsrGI, Sall |
| 5186 | CMVdel-mNeonGreen-anillin (AHD+PH) | 1645 | AgeI, BsrGI | 3331 | AgeI, BsrGI |
| 1641 | mTurquoise2-anillin(AHD+PH) | 1195 | AgeI, BsrGI | 3703 | AgeI, BsrGI |
| 5180 | CMVdel-mNeonGreen-1xaGBD(712-786) | 5186 | BsrGI, BamHI | 1st aGBD | BsrGI, BamHI |
| 5181 | CMVdel-mNeonGreen-2xaGBD(712-786/801) | 5180 | BsrGI, Aval | n+1 aGBD | BsrGI, Sall |
| 5182 | CMVdel-mNeonGreen-3xaGBD(712-786/ 801) | 5181 | BsrGI, Aval | n+1 aGBD | BsrGI, Sall |
| 5183 | CMVdel-mNeonGreen-1xpGBD(30-100)t | 5186 | BsrGI, BamHI | 1st pGBD | BsrGI, BamHI |
| 5184 | CMVdel-mNeonGreen-2xpGBD(30-100) | 5183 | BsrGI, Aval | n+1 pGBD | BsrGI, Sall |
| 5197 | CMVdel-dTomato-2xpGBD(30-100) | 5184 | AgeI, BsrGI | 4595 | AgeI, BsrGI |
| 5185 | CMVdel-mNeonGreen-3xpGBD(30-100) | 5184 | BsrGI, Aval | n+1 pGBD | BsrGI, Sall |
| 5431 | H2A-mTurquoise2-CDC42-G12V-ΔCaaX | 5068 | BsrGI, NotI | CDC42-G12V-ΔCaaX | BsrGI, NotI |
| 5432 | H2A-mTurquoise2-RAC1-G12V-ΔCaaX | 5068 | BsrGI, NotI | RAC1-G12V-ΔCaaX | BsrGI, NotI |
| 5434 | H2A-mTurquoise2-RHOA-G14V-ΔCaaX | 5068 | BsrGI, NotI | RHOA-G14V-ΔCaaX | BsrGI, NotI |
| 5712 | pLV-dTomato-2xrGBD | 5147 | EcoRV, ApaI | 5200 | EcoRV, ApaI |

To create different color fluorescent protein rGBD fusions, CMVdel-mNeonGreen-1xrGBD was digested with AgeI and BsrGI. The inserts dTomato, mScarlet-I and mTurquoise2 were digested with AgeI and BsrGI and ligated to the backbone. CMVdel-mNeonGreen-2xrGBD was digested with AgeI and BsrGI. The inserts dTomato was digested with AgeI and BsrGI and ligated to the backbone.

pLV-dT-2xrGBD was created by digesting the pLV backbone and the insert dT-2xrGBD with the restrictions enzymes EcoRV and ApaI and ligation of the two fragments.

#### aGBD

The anillin AHD + PH domain containing pEGFP-RhoA Biosensor plasmid was a gift from Michael Glotzer (Addgene plasmid # 68026). The EGFP was replaced with mTurquoise2 by making use of the AgeI and BsrGI restriction sites.

The anillin G protein Binding Domain (aGBD) was defined as the amino acid sequence from residue 712 to 786, plus a disordered linker from residue 786 to 801 (Sun et al., 2015). The aGBD was amplified by PCR with primers shown in Table 1 from the pEGFP RhoA Biosensor plasmid. The 1^st^ aGBD (residue 712-786) PCR product was digested with BsrGI and BamHI and ligated into a likewise digested CMVdel-mNeonGreen backbone, creating CMVdel-mNeonGreen-1xaGBD. For the tandem and triple aGBD the n+1 aGBD (residue 712-801) PCR product was digested with BsrGI and SalI. The backbones CMVdel-mNeonGreen-1xaGBD/ -2xaGBD were digested with BsrGI and AvaI. By ligating backbone and insert CMVdel-mNeonGreen-2xaGBD / -3xaGBD were created.

#### pGBD

The GFP-PKN1(full) was a gift from Peter Parker (Lachmann et al., 2011). The PKN1 G protein Binding Domain (pGBD) was defined as residue 30 to 100 as in the DORA biosensor (Van Unen et al., 2015). Due to its high GC content the sequence of pGBD was codon optimized with COOL (Chin et al., 2014) and the following gBlock was ordered with Integrated DNA Technologies (IDT):

5′-GA**TGTACA**GC**CTCGAG**GGGGTACAGCAGCAGCTGGAGCTGGAAAGAGAAAGGTTAAGAAGAGAGATCAGAAAAGAATTAAAGCTGAAGGAAGGAGCTGAGAACCTGAGGAGAGCCACCACAGATTTGGGAAGAAGCCTGGGCCCTGTGGAGTTATTATTAAGAGGCAGCAGCAGAAGGCTGGACCTGCTGCACCAGCAGCTGCAGGAGCTTCATGCCCAC GTGGTGCTTTAA**GGATCC**GC-3′

The pGBD was amplified from the pGBD gBlock with primers shown in Table 1. The 1^st^ pGBD PCR product was digested with BsrGI and BamHI and cloned in the likewise digested CMVdel-mNeonGreen backbone to create CMVdel-mNeonGreen-1xpGBD. For a tandem and triple pGBD the n+1 pGBD PCR product was digested with SalI and BsrGI. The backbones CMVdel-mNeonGreen-1xpGBD/ -2xpGBD were digested with BsrGI and AvaI. By ligating backbone and insert CMVdel-mNeonGreen-2xpGBD/ -3pGBD were created.

#### H2A-RhoGTPase-ΔCaaX

Cdc42-G12V, Rac1-G12V and RhoA-G14V, obtained from cDNA.org, were PCR amplified with primers shown in Table 1. To remove the CaaX motif the primer contains a mutation, replacing an A with a T, shown in bolt followed by a stop codon. The original amino acid sequence CLVL is thereby changed to S stop. The PCR products were digested with NotI and BsrGI and ligated to the likewise digested backbone H2A-mTurquiose2-N1. This results in H2A-mTurquoise2-Cdc42-G12V-ΔCaaX, H2A-mTurquoise2-Rac1-G12V-ΔCaaX and H2A-mTurquoise2-RhoA-G14V-ΔCaaX.

#### Other plasmids

The plasmid encoding Histamine 1 Receptor (H1R) was obtained from cDNA.org. The plasmids Giantin-FRB-mTurquoise2, mTurquoise2-FKBP12-p63(DH), mTurquoise2(W66A)-FRB-MoA (Addgene plasmid # 67904) and Lck-FRB-mTurquoise were described before (Van Unen et al., 2015). pLV mTurquoise2 CaaX was generated using HiFi Gibson cloning (NEB) of mTurquoise2 and CaaX into a pLV vector digested with MluI and XhoI.

### Stable Cell Line

Lentiviral particles were produced in HEK293T cells transfected with TransIT (Mirus) using 3rd generation packing plasmids (pHDMG·G VSV ENV, pHDM·HgpM2 GAG/POL, pRC-CMV-Rev1b REV) and pLV-mTurquiose2-CaaX in combination with pLV-dimericTomato-2xrGBD. Supernatant was harvested 2 and 3 days after HEK293T cell transfection, filtered (0.45 um) and concentrated using Lenti-X Concentrator (TakaraBio cat #631232). Human Umbilical Vein Endothelial Cell (HUVEC) and cord blood Blood Outgrowth Endothelial cells (cbBOEC) were transduced. Double positive cell for mTurquoise2-CaaX and dimericTomato-2xrGBD were sorted using a BD FACSAria™ cell sorter.

### Cell Culture and Sample Preparation

HeLa cells (CCL-2, American Tissue Culture Collection; Manassas, VA, USA) were cultured in Dulbecco’s modified Eagle’s medium + GlutaMAX™ (Giboc) with 10% fetal calf serum (Giboc) (DMEM + FCS) at 37°C in 7% CO2. For transfection 25 000 to 50 000 cells were seeded on round 24 mm ø coverslip (Menzel, Thermo Fisher Scientific) in a 6 well plate with 2 ml DMEM + FCS. The Transfection mix contained 1µl linear polyethylenimine (PEI, pH 7.3, Polysciences) with a concentration of 1 mg/ml per 100 ng DNA and 0.5 to 1 μg plasmid DNA per well and was mixed with 200 μl OptiMEM (Thermo Fisher Scientific) per well. After 15 min incubation at room temperature the transfection mix was added to the cells 24 h after seeding.

Blood Outgrowth Endothelial cells (BOEC) were cultivated from healthy adult donor blood as described before (Martin-Ramirez et al., 2012) and cbBOEC were cultivated from healthy donor umbilical cord. Cells were cultured in Endothelial Cell Growth Medium-2 BulletKit (CC-3162, Lonza) with 100 U/ml Penicillin (Thermo Fisher Scientific) and 100 μg/ml Streptomycin (Thermo Fisher Scientific) and 20% fetal calf serum (EGM +) at 37°C in 5% CO2. Culture dishes and coverslips were coated with 0.1% gelatin (CAS 9000-70-8, Merck) in phosphate-buffered saline 30 min prior to cell seeding. Transfection was performed with 2 μg endotoxin free plasmid DNA, using the Neon™ Electroporation Transfection system (MPK5000, Invitrogen) with the associated Neon™ Transfection System 100 μl Kit (MPK10096, Invitrogen) generating a single pulse of 1300 V for 30 ms. Cells were seeded on 24 mm ø coverslip in a 6 well plate with 2 ml EGM +.

HUVEC (Lonza,P1052, Cat #C2519A) were cultured in Endothelial Cell Growth Medium-2 BulletKit (CC-3162, Lonza) with 100 U/ml Penicillin (Thermo Fisher Scientific) and 100 μg/ml Streptomycin (Thermo Fisher Scientific) at 37°C in 5% CO2. Culture dishes and coverslips were coated with fibronectin (30 μg/mL, Sanquin) in phosphate-buffered saline 30 min prior to cell seeding.

### Spinning Disk Microscopy

Cells were imaged with a Nikon Ti-E microscope equipped with a Yokogawa CSU X-1 spinning disk unit, a 60x objective (Plan Apo VC, oil, DIC, NA 1.4), Perfect Focus System and the Nikon NIS elements software. Images were acquired with a Andor iXon 897 EMCCD camera. CFPs were imaged using a 440 nm laser line, a triple dichroic mirror (440, 514, 561 nm) and a 460 – 500 nm emission filter. GFPs were imaged using a 488 nm laser line, a triple dichroic mirror (405, 488, 561 nm) and a 500 nm long pass emission filter. RFPs were imaged using a 561 nm laser line, a triple dichroic mirror (405, 488, 561 nm) and a 600 – 660 nm emission filter. HeLa cells were imaged between 24 to 48 h after transfection in an Attofluor cell chamber (Thermo Fischer Scientific) in 1 ml of Microscopy Medium (20 mM HEPES (pH=7.4),137 mM NaCl, 5.4 mM KCl, 1.8 mM CaCl2, 0.8 mM MgCl2 and 20 mM Glucose) at 37°C. To measure the change in cytosolic intensity, HeLa cells were stimulated with 100 μM histamine and if applicable with 10 μM pyrilamine. The time trace of the cytosolic intensity of the rGBD sensor showed that the intensity stabilizes approximately 1 min after histamine addition. That allowed to compare more cells with a higher throughput by only comparing the cytosolic intensity before histamine addition to the cytosolic intensity 5 min after stimulation. For chemogenetic experiments HeLa cells were stimulated with 100 nM rapamycin (LC Laboratories).

### Widefield Microscopy

Dividing cells were imaged at a Nikon Ti-E widefield microscope, equipped with a 60x oil objective (Plan Apo λ 60x Oil), a Lumencor Spectra X light source, the Perfect Focus System, a camera (Hamamatsu C11440-22C SN:100256) and Nikon NIS elements software. HeLa cell were imaged in DMEM + FCS at 37°C and 5% CO2 in an Attofluor cell chamber in a humidified environment. RFP was imaged with an excitation wavelength of 550/15 nm and emission light was detected at 570 – 616 nm with an emission filter of 593/46 in combination with a dichroic mirror (411-452, 485-541, 567-621, 656-793 nm transmission).

### TRIF Microscopy

Cells were imaged with a Nikon Ti-E microscope equipped with a motorized TIRF Illuminator unit, a 60x TIRF objective (60x Plan Apo, Oil DIC N2, NA =1.49, WD = 120 um) and Perfect Focus System. Images were acquired with an Andor iXon 897 EMCCD camera and the Nikon NIS elements software. mTurquoise2 was imaged using the 440 nm laser line in combination with a tri split dichroic mirror (440, 488, 561 nm). DimericTomato was imaged using the 561 nm laser line in combination with a quad split dichroic mirror (405, 488, 561, 640 nm) and a dual band pass emission filter (515 to 545 nm, 600 to 650 nm). BOECs stably expressing dimericTomato-2xrGBD and mTurquoise2-CaaX were imaged in an Attofluor cell chamber in 1 ml EGM + at 37°C and 5% CO2. To measure RhoA activity in primary cells, BOECs were stimulated with 1 U/ml human α-thrombin (HCT-0020, Haematologic technologies) diluted in phosphate-buffered saline.

### Lattice light sheet microscopy

The lattice light sheet microscope located at the Advanced Imaging Center (AIC) at the Janelia Research Campus of the Howard Hughes Medical Institute (HHMI) (Chen et al., 2014) was used. HUVECs stably expressing dTomato-2xrGBD and mTurquoise2-CaaX were cultured on fibronectin-coated 5 mm round glass coverslips (Warner Instruments, Catalog # CS-5R) for 2 days. Cells were imaged at 37°C in the presence of 5% CO2 in HEPES buffer (132 mM NaCl2, 20 mM HEPES, 6 mM KCl2, 1 mM MgSO4•7H2O, 1.2 mM K2HPO4•3H2O at pH 7.4), supplemented with 1 mM CaCl2, 0.5% Albuman (Sanquin Reagents, the Netherlands), 1 g/L D-Glucose. Illumination was done using 445 nm and 560 nm diode lasers (MPB Communications) acousto-optic tunable filter (AOTF) transmittance and 100 mW initial box power and an excitation objective (Special Optics, 0.65 NA, 3.74-mm WD). Fluorescence detection was done via a detection objective (Nikon, CFI Apo LWD 25XW, 1.1 NA) and a sCMOS camera (Hamamatsu Orca Flash 4.0 v2). Point-spread functions were measured using 200 nm tetraspeck beads (Invitrogen cat# T7280) for each wavelength. Data was deskewed and deconvolved as described previously (Chen et al., 2014).

### Data Analysis

Raw microscopy images were analyzed in FIJI (Schindelin et al., 2012). Intensity time traces were generated by background correcting all images, drawing a region of interest and measuring the mean gray value for this region for each frame. To measure the cell area, a ROI was created based on a binary image generated with Huang background based thresholding for each frame in the RFP channel. The mean gray value or cell area respectively, were normalized by dividing each value by the value of the first frame. Plots for Figure 1, 2, 5 and 6 were generated with the PlotTwist web tool (Goedhart, 2020).

The change in cytosolic intensity was measured by background correcting the images, drawing a region of interest in the cytosol and measuring the mean gray value in this region. Then the ratio of mean grey value pre histamine stimulation to mean grey value post histamine stimulation was calculated and plotted as percentage using PlotsOfData (Postma & Goedhart, 2019). Scatter dot plots displaying the data points and their median were generated in this way for figure 1, 4 and 6.

To determine nuclear colocalization of the sensor with active Rho GTPases, a region of interest of the nucleus was defined by manually thresholding the signal of H2A-mTurquoise2-RhoGTPase in the CFP channel. The region of interest for the cytosol was created by enlarging the nucleus region of interest by 1 μm and subtracting the region of interest of the nucleus from this. The two regions of interest were used to measure the mean gray value of the sensor in the RFP channel in the nucleus and the cytosol. The ratio of mean nuclear intensity to mean cytosolic intensity was calculated. Plots of these nuclear to cytosolic intensity were generated for figure 3 and 4 with PlotsOfData (Postma & Goedhart, 2019).

To generate a cross section from the lattice light sheet data, slice 74 – 77 were averaged and collected in one stack.

The Kymograph was created with the Multi Kymograph function in ImageJ using a linewidth of 1 at the line indicated in the image.

## Supporting information

Supplementary information - plasmid maps

## Data availability

The following plasmids are available on addgene (http://www.addgene.org/):

\# 129633: mNeongreen-aGBD(anillin), # 129634: mNeongreen-pGBD(PKN1 codon optimized), # 129624: mNeongreen-2xrGBD, # 129625: dTomato-2xrGBD.

Sequences of the plasmids are available as supplemental information

## Author contribution

E.M. designed, analyzed and performed experiments and wrote the manuscript; F.H.v.d.L & W.v.d.M generated constructs; J.A. and J.v.B assisted with the endothelial cell experiments; S.T. performed FACS experiment; T.W.J.G. provided valuable ideas and input; J.G. designed experiments and wrote the manuscript.

## Acknowledgements

We want to thank John Heddleston and Teng-Leong Chew for the opportunity to use the Lattice Light Sheet Microscope at the Advanced Imaging Center at Janelia Research Campus. The Advanced Imaging Center at Janelia Research Campus is generously sponsored by the Howard Hughes Medical Institute and the Gordon and Betty Moore Foundation. We want to thank Ronald Breedijk for the support at the van Leeuwenhoek Centre for Advanced Microscopy, Section Molecular Cytology, Swammerdam Institute for Life Sciences, University of Amsterdam.

## Supplemental Movies

### Title and Legend for Supplemental Movies

Supplemental Movie 1. **The dimericTomato2xrGBD RhoA sensor relocating in a HeLa cell**. Spinning disk time lapse movie of a HeLa cell expressing the CMVdel-dimericTomato-2xrGBD RhoA sensor and H1R (not shown) which was stimulated with 100 μM histamine after 150 s and 10 μM pyrilamine after 350 s. The time stamper represents min:s.

Supplemental Movie 2. **The mNeonGreen1xrGBD RhoA sensor relocating in a HeLa cell**. Spinning disk time lapse movie of a HeLa cell expressing the CMVdel-mNeonGreen-1xrGBD RhoA sensor and H1R (not shown) which was stimulated with 100 μM histamine after 150 s and 10 μM pyrilamine after 350 s. The time stamper represents min:s.

Supplemental Movie 3. **Direct comparison of dimericTomato-2xrGBD sensor and mTurquise2-1xrGBD sensor in one HeLa cell**. Spinning disk time lapse movie of a HeLa cell expressing H1R (not shown) and CMVdel-dimericTomato-2xrGBD (magenta) and CMVdel-mTurquoise2-1xrGBD (grey) stimulated with 100 μM histamine. The time stamper represents min:s.

Supplemental Movie 4. **Direct comparison of dimericTomato-2xrGBD sensor and mTurquise2-AHD+PH sensor in one HeLa cell**. Spinning disk time lapse movie of a HeLa cell expressing H1R (not shown) and CMVdel-dimericTomato-2xrGBD (magenta) and mTurquoise2-AHD+PH (grey) stimulated with 100 μM histamine. The time stamper represents min:s.

Supplemental Movie 5. **AHD+PH RhoA sensor localizing in a punctate manner in an endothelial cell**. Spinning disk time lapse movie of a HUVEC expressing the eGFP-AHD+PH sensor, stimulated with 1 U/ml human α-thrombin.

Supplemental Movie 6. **Membrane specific activation of RhoA detected with the dimericTomato-2xrGBD sensor**. Spinning disk time lapse movie of a HeLa cells expressing FRB anchored to the membrane (grey), FKBP-p63-DH (not shown) and the dimericTomato-2xrGBD sensor (mpI-inferno) stimulated with 100 nM rapamycin. The time stamper represents min:s.

Supplemental Movie 7. **Mitochondria specific activation of RhoA detected with the dimericTomato-2xrGBD sensor**. Spinning disk time lapse movie of a HeLa cells expressing FRB anchored to the mitochondria (not shown), FKBP-p63-DH (grey) and the dimericTomato-2xrGBD sensor (mpI-inferno) stimulated with 100 nM rapamycin. The time stamper represents min:s.

Supplemental Movie 8. **Golgi specific activation of RhoA detected with the dimericTomato-2xrGBD sensor**. Spinning disk time lapse movie of a HeLa cells expressing FRB anchored to the Golgi (grey), FKBP-p63-DH (not shown) and the dimericTomato-2xrGBD sensor (mpI-inferno) stimulated with 100 nM rapamycin. The time stamper represents min:s.

Supplemental Movie 9. **Visualization of active RhoA at the cleavage furrow with the dimericTomato-2xrGBD sensor**. Wide field time lapse movie of a HeLa cell expressing dimericTomato-2xrGBD. The time stamper represents min:s. LUT = mpI-inferno.

Supplemental Movie 10. **Visualization of active RhoA at the contracting cell edge in a randomly migrating endothelial cell**. Spinning disk time lapse movie of a BOEC expressing the dimericTomato-2xrGBD sensor. The time stamper represents min:s. LUT = mpI-inferno.

Supplemental Movie 11. **Visualization of active RhoA at the basolateral plasma membrane of an endothelial cell**. TRIF microscopy time lapse movie of a BOEC stably expressing the dimericTomato-2xrGBD sensor (left, LUT = mpl-magma) and mTq2-CaaX (middle, grey), stimulated with 1 U/ml human α-thrombin. The right shows a binary image representing the cell area. The time stamper represents min:s.

Supplemental Movie 12. **Visualization of active RhoA at the plasma membrane in a cross section**. Cross section of a lattice light sheet time lapse movie of a HUVEC stably expressing dimericTomato-2xrGBD sensor (upper panel, LUT = mpl-magma) and mTq2-CaaX (lower panel, grey), stimulated with 2,5 µM nocodazole 10 min prior to the imaging.

Supplemental Movie 13. **Whole field of view for cross section of Supplemental Movie 12**. Maximum Intensity projection for the whole field of view of the cell that the cross section is presented for supplemental movie 12. Lattice light sheet time lapse movie of a HUVEC expressing dimericTomato-2xrGBD sensor (upper panel, LUT = mpl-magma) and mTq2-CaaX (lower panel, grey), stimulated with 2,5 µM nocodazole 10 min prior to the imaging. The cross section was taken vertically at three quarters from the left edge of the image.

Supplemental Movie 14. **Visualization of active RhoA in a contracting endothelial cell**. TIRF microscopy time lapse movie of a BOEC stably expressing the dimericTomato-2xrGBD sensor (left, LUT = mpl-magma) and mTq2-CaaX (right, grey) stimulated with 1 U/ml human α-thrombin 5 min prior imaging. The time stamper represents min:s.

